# ViCloD, an interactive web tool for visualizing B cell repertoires and analyzing intraclonal diversities: application to human B-cell tumors

**DOI:** 10.1101/2022.11.28.518196

**Authors:** Lucile Jeusset, Nika Abdollahi, Thibaud Verny, Marine Armand, Anne Langlois De Septenville, Frédéric Davi, Juliana S. Bernardes

**Affiliations:** Sorbonne Université, CNRS, UMR 7238, Laboratoire de Biologie Computationnelle et Quantitative, Paris, France; Sorbonne Université, AP-HP, Hôpital Pitié-Salpêtrière, Department of Biological Hematology, Paris, France; Ecole des Mines ParisTech, Paris, France; IMGT^®^, the international ImMunoGeneTics Information System, CNRS, Institute of Human Genetics, Montpellier University, France

**Keywords:** B-cell immune repertoire, intraclonal diversity, B cell lineage trees, immunoinformatics

## Abstract

High throughput sequencing of adaptive immune receptor repertoire (AIRR-seq) has provided numerous human immunoglobulin (IG) sequences allowing specific B cell receptor (BCR) studies such as the antigen-driven evolution of antibodies (soluble forms of the membrane-bound IG part of the BCR). AIRR-seq data allows researchers to examine intraclonal differences caused primarily by somatic hypermutations in IG genes and affinity maturation. Exploring this essential adaptive immunity process could help elucidate the generation of antibodies with high affinity or broadly neutralizing activities. Retracing their evolutionary history could also help to clarify how vaccines or pathogen exposition drive the humoral immune response. Computational methods are necessary for large-scale analysis of AIRR-seq properties. However, there is no efficient and interactive tool for analyzing intraclonal diversity, permitting users to explore adaptive immune receptor repertoires in biological and clinical applications. Here we developed ViCloD, a web server for large-scale visual analysis of repertoire clonality and intraclonal diversity. ViCloD uses data preprocessed by IMGT/HighV-QUEST and performs clonal grouping and evolutionary analyses, producing a collection of useful plots. The web server presents diverse functionalities, including repertoire navigation, clonal abundance analysis, and intraclonal evolutionary tree reconstruction. Users can download the analyzed data in different table formats and save the generated plots as images. ViCloD is a simple, versatile, and user-friendly tool that can help researchers and clinicians to analyze B cell intraclonal diversity. Moreover, its pipeline is optimized to process hundreds of thousands of sequences within a few minutes, allowing an efficient investigation of large and complex repertoires.

**Availability and implementation:** The web server is available at http://www.lcqb.upmc.fr/viclod/. The pipeline is available at github and allows users to run analyses locally https://github.com/julibinho/ViCLoD

## 1 Introduction

B cells are critical components of the adaptive immune system, recognizing and eliminating various foreign molecules (antigens). B cells recognize antigens through a specific protein complex, the B cell Receptor (BCR), attached to the cell surface. The BCR is made up of a recognition unit formed by an immunoglobulin (IG) protein and an associated signaling unit, the Ig*α* and Ig*β* molecules. In humans, the IG consists of four polypeptide chains: two heavy chains and two light chains, joined by disulfide bonds. Besides the membrane-bound form, IG can also be secreted in the blood plasma as circulating antibodies.

The immune system can respond to almost any antigen to which it is exposed, partially through the vast diversity of BCRs expressed by B cells. Such diversity lies in the variable region of IG gene and results from complex genetic mechanisms that occur during B cell ontogeny. This process, called VDJ recombination (*1*), joins different types of genes: V (variable), D (diversity) in heavy chain, and J (joining). The resulting combinatorial diversity is further enhanced by a junctional diversity due to the imprecise joining of these genes with deletion and insertion of random nucleotides at the recombination sites, thereby creating an extremely diverse zone within the variable region, the complementary determining region 3 (CDR3) (*2*). Further diversity occurs after the antigen encounter, with a process called affinity maturation, which occurs in specialized structures of secondary lymphoid organs (*3*). There, point mutations are introduced in the variable region of IG genes at very high rates, about 10^6^ times higher than observed for other genes, by a mechanism termed somatic hypermutation (SHM) (*4*). This is followed by a Darwinian selection process whereby B cells expressing BCR with higher affinity resulting from SHM survive and proliferate in the germinal centers (*5*). Therefore, the generation and selection of somatic antibody mutants are decisive for increasing antigen affinity, contributing to the B cell immune repertoire diversity and plays a fundamental role in an efficient immune response (*6*).

The adaptive immune receptor repertoire sequencing (AIRR-seq) has intensively contributed to characterizing immune repertoires, potentially revealing the high-dimensional complexity of the immune receptor sequence landscape. AIRR-seq data is often used to address many scientific and clinical questions. For instance, an extremely diversified BCR repertoire reflects the expected diversity in healthy individuals. However, such diversity can be significantly affected by different factors such as autoimmune diseases (*7*–*13*), allergy (*14*–*16*), cancer (*17*, *18*), and aging (*19*, *20*). The exploration of AIRR-seq data is thus an active field of research, mainly to analyze BCR composition and diversification. Analyzing mutational sequence patterns of a given clonal lineage can provide insights into the diversification and selection processes that lead to B cell clonal evolution. The examination of intraclonal compositions can also help in understanding the response to infections (*21*), vaccines (*22*, *23*), the mechanisms of autoimmune diseases (*24*) and allergy (*25*), among others. Rapid advances in high-throughput sequencing have opened up new possibilities for studying B cell clonal expansion and intraclonal compositions. With decreasing costs of DNA sequencing technology, numerous IG sequences with inherent complexity, variability, and mutational load are accessible, including in clinical contexts. Thus, the availability of AIRR-seq data has motivated researchers with different backgrounds (biological, computational, and statistical) to investigate and examine the adaptive immune complexity.

Several methods and computational tools have been developed to treat different AIRR-seq analysis steps, producing multiple integrated context-specific software (*26*–*29*). Most of the tools were designed for V(D)J assignments (*30*), or the identification of groups of clonally-related sequences (*31*–*33*). Some solutions exist to tackle and determine the heterogeneity of BCR repertoires (*29*, *34*–*37*). Such tools quantify the diversity of a BCR repertoire by estimating clonal group sizes, characterizing CDR3 regions, identifying the most used V, D, and J genes (alleles), and the most frequent VJ combinations. However, few tools allow the investigation of intraclonal diversity/evolution of a B-cell repertoire (*38*–*42*). Such analyses can potentially reconstruct antibodies’ evolutionary (clonal) development and identify mutable positions that correlate with antigen-binding sites. Most existing methods are stand-alone programs requiring computational skills for installation and pre-processing tasks such as clonal grouping and lineage tree inferences. Furthermore, such tools do not provide interactive plots that could assist users in their analyzes. Thus, we are currently lacking a large-scale, time-efficient and versatile AIRR-seq tool for deeply analyzing intraclonal diversity and evolution.

Here we propose ViCloD, a web-based interactive tool for visualizing B cell receptor repertoire clonality and analyzing intraclonal diversity. ViCloD enables repertoire analysis and proposes a dynamic repertoire topology panorama. Users can navigate among B-cell clonal lineages and explore their intraclonal diversity. For this purpose, we have developed a set of algorithms to analyze millions of IG sequences within a few minutes (*31*, *43*). Such methods are time efficient and compatible with both research and clinical settings. Users can download all processed data, save the generated plots, perform their analyses locally, and visualize them in the ViCloD web server. our web-server is a user-friendly data visualization tool to help the research/medical community select helpful information in BCR repertoire analyses. It allows non-computational experts to analyze intraclonal diversities, boosting their research within the immune repertoire investigation. ViCloD provides unique functionalities absent in other web servers dedicated to AIRR data processing, analysis, and visualization. It is a versatile AIRR-seq tool with the potential to help experimental/clinical investigations of immune repertoires. To demonstrate its functionalities and potential, we applied it to repertoires of human B-cell tumors. However, ViCloD can be used to investigate B-cell clonal expansions and intraclonal composition of any type of repertoire.

## 2 Material and Methods

### 2.1 ViCloD workflow

The ViCloD workflow is presented in Figure 1; it consists of five main steps: upload input, data processing (pipeline), repertoire composition, clonal composition, and lineage architecture, described below.

**Figure 1:**
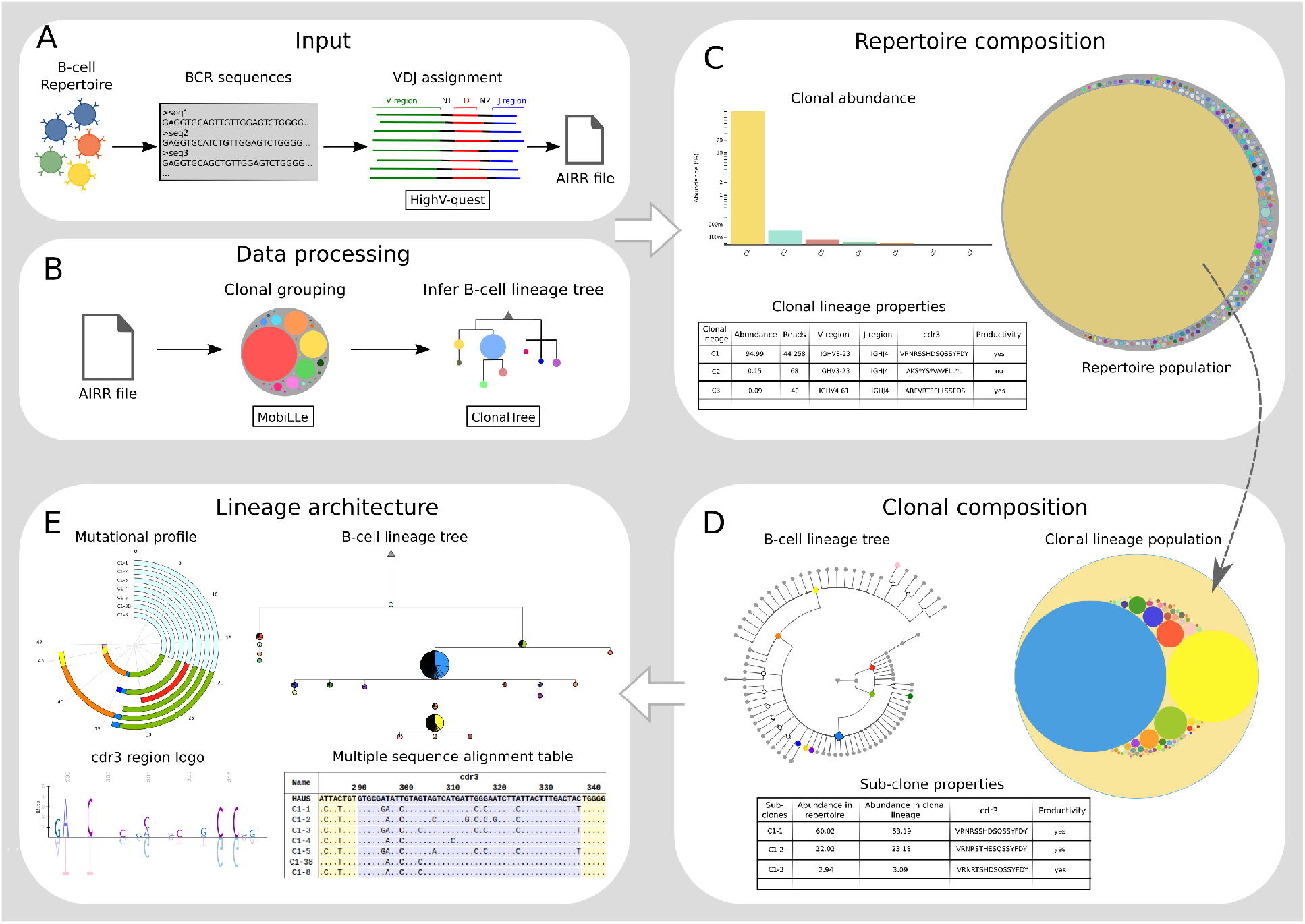
ViCloD Flowchart. (A) Input. IGH sequences of a given repertoire are previously annotated and uploaded within ViCloD web server as an AIRR file. (B) Data processing. MobiLLe processes the AIRR file and infers B-cell clonal lineages, which are then analyzed by clonalTree to reconstruct their evolutionary tree. (C) Repertoire composition. Clonal lineages are represented as nested circles, a bar plot shows the abundance of the most predominant, and a table summarizes some of their essential properties. (D) Clonal composition. Users can explore clonal composition by selecting a B-cell clonal lineage. Circles represent subclones, and a circular B-cell lineage tree represents the evolutionary relationships. A table shows additional properties of subclones. (E) Lineage architecture. Users can explore a B-cell lineage inferred tree by an interactive tree, a circular bar chart (mutational profile), a CDR3 logo, and multiple sequence alignments.

#### 2.1.1 Uploading input

As IGH captures most of intraclonal diversification, ViCloD has been evaluated so far only on IGH repertoires. IGH sequences should be first filtered upon quality control and annotated to determine their IGHV, IGHD, IGHJ genes and localize their CDR3 region. ViCloD accepts data from IMGT/HighV-QUEST (*44*), the international standard web software for annotating IG sequences, see Figure 1-A. In principle, the output of any V(D)J annotation tool could be used as long as it provides the minimum information needed for the analysis, specified in http://www.lcqb.upmc.fr/viclod/Format. ViCloD requires AIRR formatted input files (*45*) that facilitate using other V(D)J assignment tools rather than IMGT/HighV-QUEST. Users can upload the AIRR file on the main page of the ViCloD web server, enter their email address to receive a link to their outputs, and set a threshold *t* for eliminating infrequent reads; *t* can have an absolute or percentage value. The analysis is relatively quick; for instance, it can take around two minutes to analyze a monoclonal repertoire containing around 300K sequences. However, the computational time can vary depending on the structure of the sample (the size/amount of clonal lineages and sequence mutations); see some examples in Table S1. Users can also submit files previously processed by ViCloD: downloaded from the web server, or generated by the stand-alone version. Thus, users can store and re-analyze their data without any time constraints.

#### 2.1.2 Data processing

Once IGH sequences are uploaded, the first step is to group them into clonally-related sequences. We use MobiLLe (*31*), a multi-objective-based clustering for inferring B-cell clonal lineages from high-throughput AIRR-seq data. Within each B-cell clonal lineage detected by MobiLLe, we group sequences with the same IGHV, IGHJ genes, and identical CDR3 amino acid sequences, forming *subclones*. Note that we adopted the IMGT definition (*46*) to define a subclone, whereas we consider a clonotype a group of identical nucleotide sequences, see Figure S1. For each clonal lineage, we try to reconstruct its evolutionary tree by using the *N* most abundant subclones. We use ClonalTree (*43*), a time-efficient and accurate approach to reconstruct B-cell lineage trees that incorporates genotype abundances into minimum spanning tree algorithms. ClonalTree requires a file in FASTA format containing sequences and their abundances as input. Each sequence in the input file represents a subclone. Since sequences in a given subclone can differ due to SHM, we select the most abundant sequence to represent it. The abundance corresponds to the total number of sequences within a subclone. Clonal-Tree also requires the hypothetical ancestral unmutated sequence (HAUS) representing the tree root.

For that purpose, we consider the IGHV and IGHJ germline sequences provided by IMGT/HighV-QUEST. For the junction part, we select the sequence with the lowest number of mutations on the IGHV gene compared to the germline, then we extract its junction region. Finally, we concatenate the three parts (IGHV, IGHJ, and junction region) to obtain the HAUS. The Data processing pipeline is illustrated in Figure 1-B.

#### 2.1.3 Repertoire composition

We used nested circles to represent B-cell clonal lineages within a repertoire, Figure 1-C. The outer circle represents the entire BCR repertoire, while internal circles correspond to B-cell clonal lineages. The size of each circle correlates with the clonal lineages’ abundance. A bar plot on the same web page shows the abundance of the ten most predominant clonal lineages. Users can choose between a standard or logarithmic scale; they can also select a threshold of clonal lineage abundance when analyzing the clonality of a repertoire, for instance, above 5%. When such a threshold is set, only B-cell clonal lineages with an abundance higher than this value are retained in the BCR repertoire visualization; all others are hidden. A table at the bottom of the page summarizes some essential properties of each inferred B-cell clonal lineages, such as its identifier, its abundance, the IGHV/IGHJ gene annotations, the CDR3 amino acid sequence and the “functionality” status of the most abundant clonal lineage sequence, where ‘yes’ indicates productive sequences, and ‘no’ unproductive ones.

#### 2.1.4 Clonal composition

Users can explore clonal composition by clicking on one of the circles representing a B-cell clonal lineage. The circle expands and other circles, representing subclones, appear (see dotted arrow in Figure 1-CD). Note that we label only the subclones with abundance higher than 1%. The circle area correlates with the subclone abundance within the B-cell clonal lineage; see panel “Clonal lineage population” in Figure 1-D. For the five most abundant B cell lineages, a circular B-cell lineage tree generated by ClonalTree is displayed, representing the evolutionary relationships among such sub-clones and the HAUS. Several options for tree representation are available. For instance, nodes can be colored by subclones, a different color for each one, or by functionality, where green and red nodes represent productive and unproductive rearrangements, respectively. The latter is due to stop codons or out-of-frame IGHV-IGHD-IGHJ junctions. It is also possible to display the length of tree branches representing distances between sequences. The HAUS is represented by a triangle and the most abundant subclone by a square to facilitate tree interpretation. At the bottom of the page, additional information about subclones, such as their identifiers, abundances, and CDR3 amino acid sequences, is displayed.

The lineage tree inferred by the ClonalTree algorithm can represent (at most) 200 of the most abundant subclones. Interpreting the complete lineage tree can be challenging for large clonal lineages containing many subclones. Thus, we propose two simplified tree topologies while conserving nodes with high abundance and their path from the root. We developed two strategies for pruning the tree (*47*) for a more comprehensible visualization, Figure S2. These simplified versions may be more valuable than an entire tree with many details. The first pruning strategy transforms the tree shown into Figure S2-A in the tree of Figure S2-B; it eliminates nodes with no descendant or an abundance lower than a predefined threshold (by default 0.01%). This approach is more helpful for trees containing few nodes. Trees with many subclones can require the second pruning strategy that eliminates nodes with low abundance if they present high similarity with more abundant subclones (Figure S2-BC). For both strategies, we eliminate just leaf nodes and keep the *N* most abundant subclones in the simplified tree (by default, N=10). We try to preserve the tree’s evolutionary paths by eliminating only leaves’ nodes, which could be destroyed if an internal node is removed. We keep the *N* largest subclones since they contain relevant information to analyze the lineage tree. We apply the first strategy once, and on the resulting tree, we repeat the second one several times until achieving at most 30 nodes. Users can navigate between the complete lineage tree and pruned trees on the website.

#### 2.1.5 Lineage architecture

The last ViCloD module provides more details about the B-cell lineage inferred tree, Figure 1-E. Users can explore the simplest tree to examine intraclonal diversities. ViCloD provides four types of representations: an interactive tree, a circular bar chart (mutational profile), a CDR3 logo, and a multiple sequence alignment.

##### Interactive lineage tree

We kept the most pruned tree with at least N=10 most abundant subclones for an easier exploration. The coloured circles indicate observed subclones, while the white circles represent unobserved nodes, which reflect unobserved sequences that were not sampled or eliminated during affinity maturation (cf Clonaltree algorithm). The HAUS is represented by a triangle. The branch length depicts the number of SHM between the representative sequences of connected subclones or the HAUS. We designate subclones by a number associated with their abundance, so the subclone 1 is the most abundant in the B-cell lineage. In the tree, nodes representing the five largest subclones have a bold border and are numbered from 1 to 5. When the user moves the cursor over nodes, subclone details (identifier, abundance, and functionality) appear. The option “display abundance” displays each subclone abundance by increasing the node’s diameter. In this visualization mode, nodes are labeled with their abundance (%) rather than with a sequential number. Each subclone may contain multiple unique sequences (clonotypes). Choosing “Display proportion of clonotypes” will turn each node into a pie chart, illustrating the numerical proportion of unique sequences within the subclone. A pie chart for each node can be viewed on a separate page by right-clicking on the node and selecting “See the distribution of clonotypes” (Figure S3).

##### Circular bar chart - Mutational profile

The circular bar chart, also termed as “Mutational profile”, shows the cumulative path distances for a set of selected subclones. By default, ViCloD displays the five most abundant subclones, but any subclone can be included and/or removed from the plot; a maximum of eight items are displayed. Each colored section is related to a subclone and represents the number of point mutations observed between a given subclone and its parent in the tree. To highlight the tree path from a subclone to the root, users need to hover the mouse over the desired subclone identifiers. To highlight a branch in the tree, one can hover the mouse over each section of the circular bar chart. The number of mutations between the pair of connected subclones are also displayed.

##### Multiple sequence alignment

To illustrate the conservation/mutations between the representative sequence of each subclone and the HAUS, we build a multiple sequence alignment (MSA) with the MUSCLE program (*48*). For each subclone, we display in a table: its identifier, the percentage and number of reads, the divergence rate (number of mutations) between the subclone representative sequence and the HAUS, and the percent identity of IGHV sequence when compared to the HAUS. Only mutated nucleotides of the subclone representative sequence are shown in the alignment, whereas a dot represents conserved positions. CDR and framework sections are highlighted with different colors, and the IGHD gene is underlined in each sequence; see an example in Figure 2. Users can sort out sequences in the table based on each column. The table can be downloaded into a CSV file by clicking on the download button. It is also possible to select one, multiple, or all sequences and send them directly to the IMGT/HighV-quest web-server. A button on the top-right allows users to visualise a simplified MSA that can make its interpretability easier. A second button displays the complete alignment containing all subclones.

**Figure 2:**
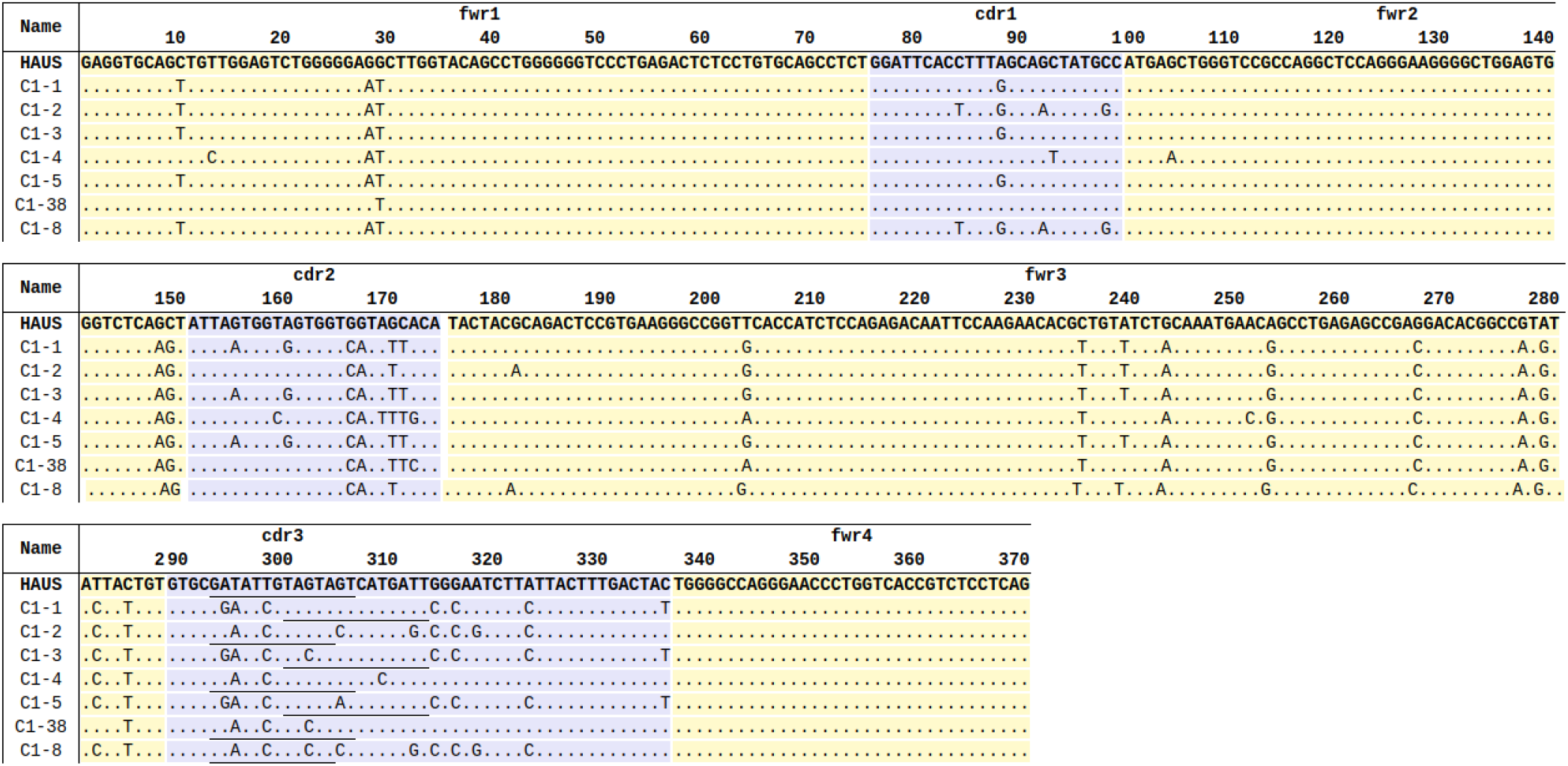
Multiple sequence alignment of representative sequences of clonal lineage C1 (Patient. **1)** Sequence alignment of seven subclones of clonal lineage C1: the five most abundant, the closest (C1-38) and the most distant (C1-8) to the HAUS.

##### CDR3 logo

Sequence logos provide a more detailed and precise description of sequence similarities and are frequently used to unravel significant features difficult to visualize in the multiple sequence alignment. In the “Lineage architecture” page, users can generate a sequence logo of the CDR3 region by clicking on the “CDR3 Logo” button. The logo is split into two sections; the upper one shows the most frequent amino acids that differ from the HAUS, while the lower section shows the corresponding residues in the HAUS. The height of the stack letter represents the frequency of each residue in the alignment, showing the mutational burden in the clonal lineage; see an example in Figure 3.

**Figure 3:**
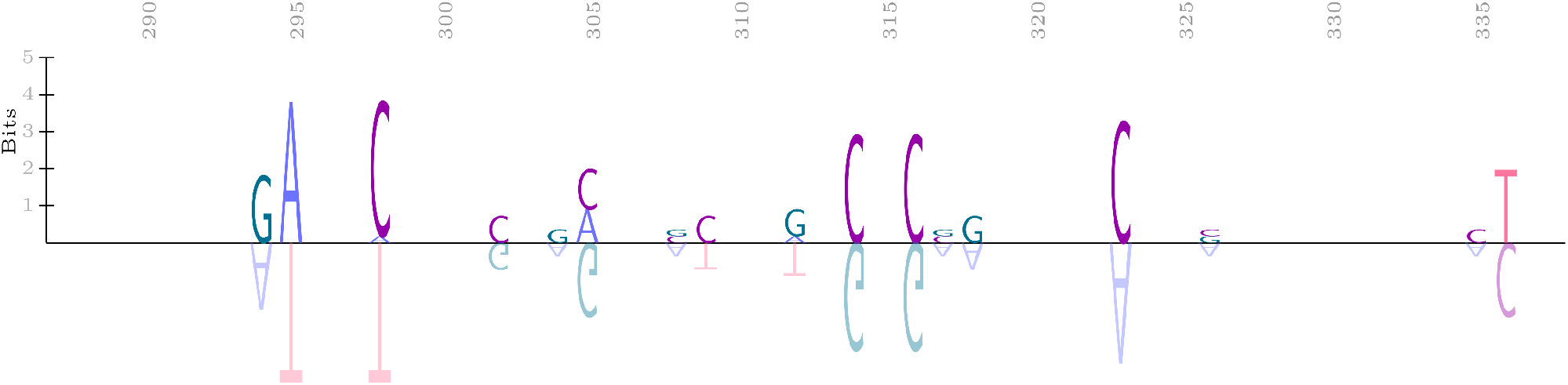
CDR3 logo. The upper section depicts the most frequent amino acids that differ from the HAUS, while the lower section shows the corresponding HAUS residues.

### 2.2 Implementation

The server side of ViCloD website was implemented in python (version 3), Bash script and PHP (version 7.4.3, http://php.net/) that was used for interactive data processing. We also used the MySql database (mysql.com/) to control the user’s pending jobs. The client-side web interface was implemented using HTML5, CSS, and JavaScript (javascript.com/) along with several libraries, including D3.js (d3js.org/), FileSaver.js (github.com/eligrey/FileSaver.js/), saveSVGAsPNG (github.com/exupero/saveSvgAsPng), PDFKit (pdfkit.org/), and JSZip (stuk.github.io/jszip/). Users submit their input files and can provide an email address to receive their results. We first check that the input file contains all the required information. Then, data are processed by the ViCloD Data processing tools, and all analyses are directly available or by clicking on the link sent by email. Users’ files (inputs and outputs) are available for download (for ten days) and can be retrieved during this time. Users can also download JSON files (json.org) that can be afterwards uploaded into the ViCloD web server. It enables data to be stored locally and re-analyzed without time constraints.

### 2.3 Tutorials

The “Tutorial” section of the website includes a variety of text-based tutorials to help users getting started with ViCloD. Multiple analyzed data sets are provided on the “Example” page to assist users step by step. Several sections under the “Overview tab” provide additional information, such as references, algorithms used, and the server’s versions.

### 2.4 Experimental data

IGH-VDJ rearrangement sequences were obtained from patients with B cell malignancies during standard routine diagnostic or prognostic procedures at the Pitié-Salpêtrière Hospital. Informed consent was obtained from patients according to local regulations. Genomic DNA was extracted from peripheral blood or tissue biopsy, and IGH-VDJ next-generation sequencing was performed as previously published (*49*). Four patients with either follicular lymphoma or chronic lymphocytic leukemia were selected based on the findings of intraclonal diversity in their tumor clonal IGH-VDJ gene rearrangements.

### 2.5 Data Availability

ViCloD web-server is available at http://www.lcqb.upmc.fr/viclod/, where one can also find user manual and repertoire examples. The stand-a-lone version is available at https://github.com/julibinho/ViCLoD.

## 3 Results

BCR immune repertoire studies have a significant impact in a variety of research and clinical areas. Several works have reported a wide range of intraclonal diversity when analyzing BCR repertoires of patients with different B-cell malignancies (*50*–*52*). In the following section, we applied ViCloD to BCR repertoires from two B-cell malignancies, illustrating different levels of intraclonal diversities.

### 3.1 Clonal heterogeneity in a case of follicular lymphoma

Follicular lymphoma (FL) is one of the most frequent types of non-Hodgkin lymphomas, and is often a relatively indolent disease, although it can transform into an aggressive form. It is a tumor of mature B lymphocytes, typically developing in lymph nodes, and originates from germinal centers, where most of SHM occur. Hence, lymphomatous FL cells express surface IG, which carry high levels of SHM in their IG variable regions, comparable with that of normal germinal center B cells (*50*). The process of SHM may continue in lymphomatous cells leading to clonal evolution and emergence of subclones over time (*53*, *54*).

We used the IGH repertoire of a patient with FL (labeled patient 1) to demonstrate all types of analyses provided by ViCloD, see http://www.lcqb.upmc.fr/viclod/Examples/example1. Around 186K sequences from the patient were uploaded, but only 46K (≈ 25%) were analyzed after quality control and filtering (*t* = 0, 005%); see Section 2.1.1. Figure 4-A shows the repertoire composition of patient 1, with the presence of dominant clonal lineage accounting for more than 94% of the repertoire, annotated with IGHV3-23 and IGHJ4 genes. Such clonal proliferation is expected in this type of B-cell malignancy. Minor unrelated clonal lineages, each at a frequency below 0.2% were also detected, probably originating from the non-tumoral polyclonal background of B cells, see Table S2. ViCloD allows users to examine the five most abundant clonal proliferation, labeled in Figure 4-A.

**Figure 4:**
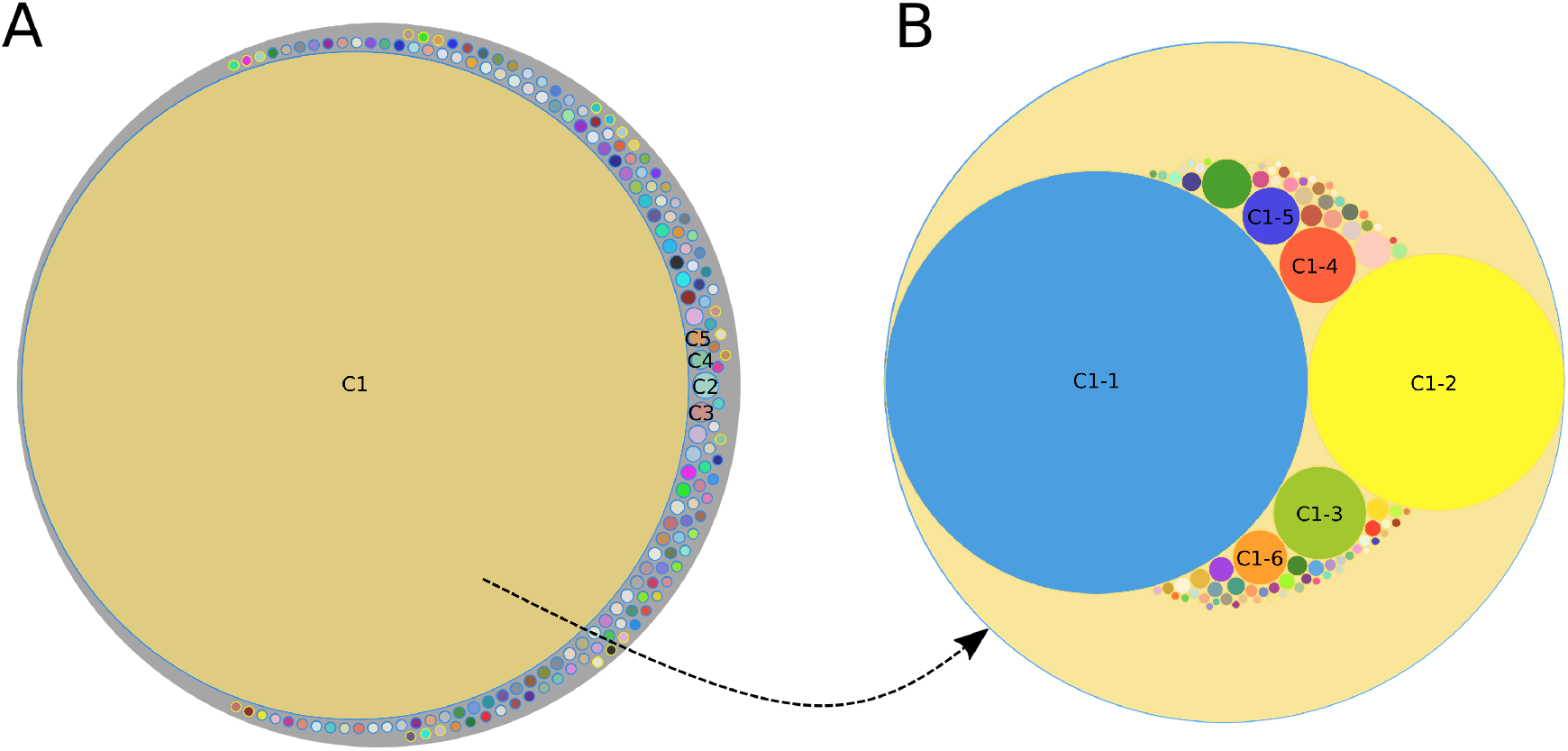
Repertoire composition of a patient with follicular lymphoma (Patient 1). (A) circles represent the most abundant clonal lineages; Circle area correlates with clonal lineage abundance. (B) Clonal population of the most predominant clonal lineage (C1). Circle area correlates with subclone abundance.

Users can click on a given clonal lineage to inspect its composition, see Section 2.1.4. Figure 4-B shows subclones for the most predominant clonal lineage C1. For this example, we labeled only six subclones, with an abundance greater than 1%, but 87 were detected, indicating a high level of intraclonal diversity for C1 lineage. Sequences of all subclones are productive. Subclones C1-1 and C1-2 are the most abundant of the C1 lineage, grouping around 60% and 22% of IGH sequences, respectively. We also observed intraclonal diversity for other clonal lineages; more details are available on the “Examples tab” at ViCloD website, see the URL above.

ViCloD also provides lineage architecture details for a given clonal lineage. Figure 5 shows the B-cell lineage tree and mutational profile for the clonal lineage C1. In the tree, Figure 5-A, the HAUS is indicated by a triangle, while the most relevant subclones appear as circles. The circle area is proportional to the sub-clonal abundance, and pie charts indicate the sub-clonal composition with the proportion of clonotypes. For instance, the subclone C1-1 is composed by 536 different clonotypes, while C1-2 by 255; see clonotype distributions for the five most abundant subclones in Figure S3. The tree topology confirms the high intraclonal diversity of lineage C1. The reconstructed tree has several levels, revealing the intricate evolutionary relationships of the most abundant/relevant subclones. Accordingly, subclone C1-1 gave rise to seven subclones, and C1-2 evolved from a minor subclone, C1-6. At the top of the tree, subclone C1-38, which accounts for less than 0.05% of the clonal lineage C1, is the most conserved, with a lower mutation rate (19 nucleotide differences) compared to the HAUS. The longest path in the tree is until subclones C1-8, C1-30, and C1-39, which group 0.49%, 0.07%, and 0.05% of the lineage, respectively, and harbor the highest number of mutations (n = 47). Interestingly, the subclone C1-4 presents several descendants, but none of them is among the most abundant.

**Figure 5:**
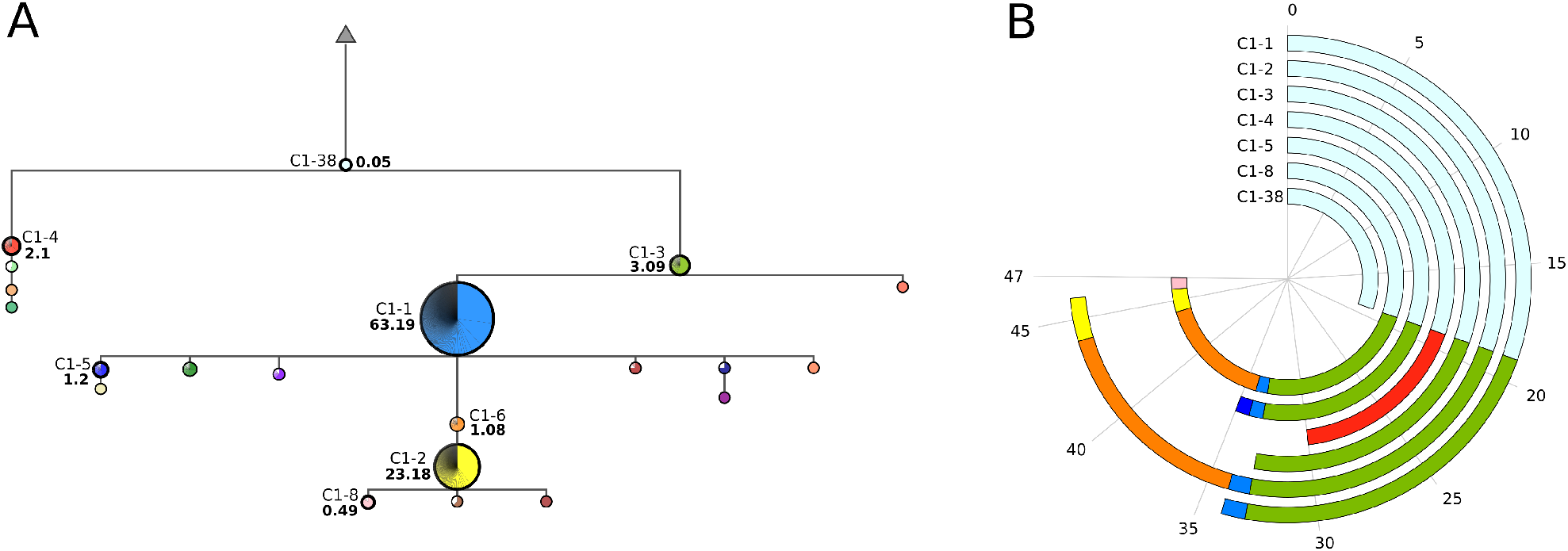
Lineage architecture (Patient 1). (A) simplified version of the B-cell lineage tree. (B) cumulative path distance between the HAUS and the selected subclones.

Figure 5-B shows a circular plot with cumulative distances for the seven most abundant/relevant subclones of the C1 lineage. Each colored arc represents a subclone, and the arc length corresponds to the number of mutations for its direct parent. The number of cumulative mutations (CM) of a given subclone is the sum of all arc lengths and corresponds to the path length from the tree root to a specific subclone. For instance, we observe 47 cumulative mutations from C1-8 and only 19 for C1-38, as can be seen when comparing arc lengths of C1-8 and C1-38 in Figure 5-B. The circular plot (mutational profile) is interactive, allowing users to add/remove subclones by clicking on nodes in the tree; a maximum of eight subclones can be represented simultaneously. Moving the mouse over a subclone on the circular plot shows its evolutionary path until the root (highlighted in red in the tree).

Cumulative mutations can be further examined in Figures 2 and 3: the former shows multiple sequence alignment (MSA) along the entire IGH-VDJ rearrangement, while the later displays a sequence logo centered on the CDR3 region. Parts of sequences in MSA are colored according to their structural framework regions (FR, yellow) and complementary determining regions (CDR, violet). Conserved positions are indicated by dots, while the presence of a nucleotide indicates a mutation in that position. The IGHD gene is underlined within the CDR3, which is the most diverse part of IGH sequences. The sequence logo helps examine mutations in that region, Figure 2.

### 3.2 ViCloD identifies different types of clonal lineage architectures

Chronic lymphocytic leukemia (CLL) is another frequent B-cell malignancy, characterized by accumulation in the blood of mature B clonal B cells. Sequencing of IG expressed by the leukemic cells has revealed two subgroups of CLL depending on the mutation load of the IGHV genes. In addition, this parameter constitutes a strong prognostic indicator as patients with no or few mutations have a more aggressive disease (*51*). Recent high-throughput IGH repertoire sequencing experiments has revealed a much higher intraclonal diversity than anticipated, although the exact mechanisms leading to this ongoing SHM activity remains elusive (*52*). We selected three patients with CLL to demonstrate how ViCloD can analyze repertoires having different levels of intraclonal diversity. We selected the major clonal lineage from the repertoire of patients 2, 3 and 4 that correspond to 96%, 99% and 99% of the repertoire, respectively.

Figure 6 shows B-cell lineage tree topologies and mutational profiles for the most abundant clonal lineages of selected individuals. When comparing the clonal trees of the three patients (Figure 6-A,C,E), we observe different topologies. The clonal tree of patient 2 is very branched; the most abundant subclone C1-1 descends from C1-2, the second most abundant. C1-2 presented eight cumulative mutations, while C1-1 14; we observe six point mutations between them. We can also see the high intraclonal diversity of patient 2 in the mutational profile shown in Figure 6-B. Subclone C1-33 is the closest to the HAUS with seven nucleotide differences, while C1-51 has the most mutated sequence with 27 cumulative mutations. The third most abundant subclone, C1-3, is also highly mutated with 22 cumulative mutations. C1-3 descends from a less abundant subclone C1-5 that originates from C1-6.

**Figure 6:**
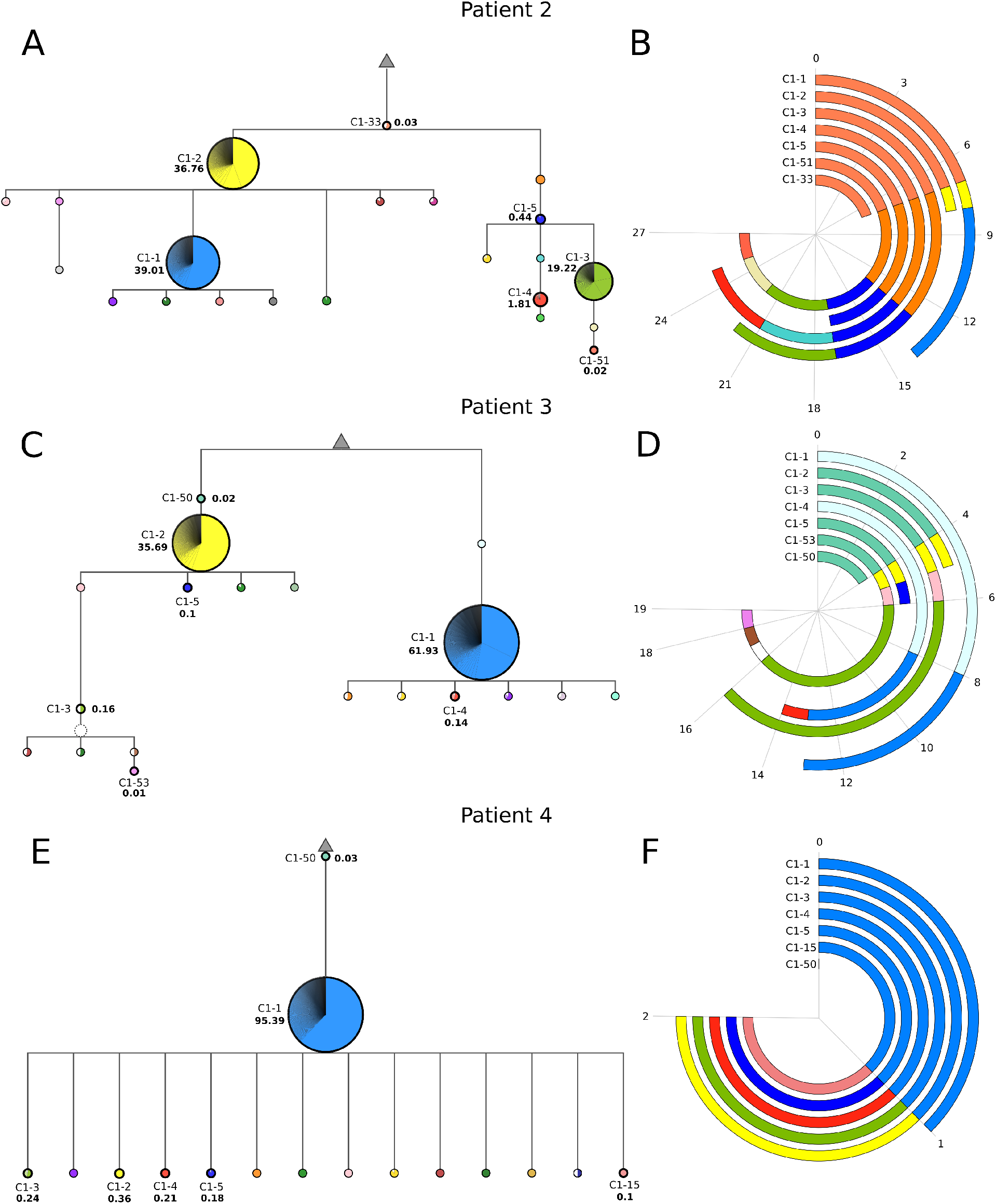
Clonal lineage architectures for three patients with CLL. We show the simplest B-cell lineage tree (A, C, E) and the mutational profile (B, D, F) for patients 2, 3 and 4, respectively. Mutational profiles contain seven subclones: the five most abundant, the closest and the furthest to HAUS.

In patient 3, we also observed some intraclonal diversity, but with a different tree topology and mutational profile, when compared to patient 2; compare Figure 6-AB and Figure 6-CD. The most abundant subclone of patient 3 (C1-1) grouped 61.93% of clonal lineage sequences and has 13 cumulative mutations. subclone C1-2, with an abundance of 35.69%, presented five cumulative mutations; between C1-1 and C1-2 we observed ten nucleotide differences. Both subclones share a few point mutations, which is why the ClonalTree algorithm puts them in different tree branches. The intraclonal diversity is also evident in the mutational profile of patient 3, Figure 6-D shows a minimum of four (C1-50) and a maximum of 19 (C1-53) cumulative mutations.

On the other hand, patient 4 shows lower intraclonal diversity, which is clearly observed in its tree topology and mutational profile, Figure 6-EF. This patient’s repertoire has one predominant subclone (C1-1), representing more than 95% of both clonal lineage and entire repertoire. It has just one cumulative mutation, while the subclone C1-50 has zero. We also observed a tree with few levels (only three), see Figure 6-E. A total of 86 subclones were detected. All subclones (except C1-50) descend from C1-1, and all C1-1 descendants present just two cumulative mutations, and thus, just one more than C1-1. Note that all mutations were observed in the CDR3 regions, and interestingly, they involve predominantly ‘G’ and to a lesser extent ‘C’ nucleotides.

### 3.3 Comparison with other clonal lineage visualization tools

To compare the functionalities of ViCloD with other existing methods, we chose four AIRR-seq tools, which provide both evolutionary analysis (clonal lineage tree/graph) and intraclonal visualizations. Table 1 shows the main differences between ViCloD and the considered tools.

**Table 1:**
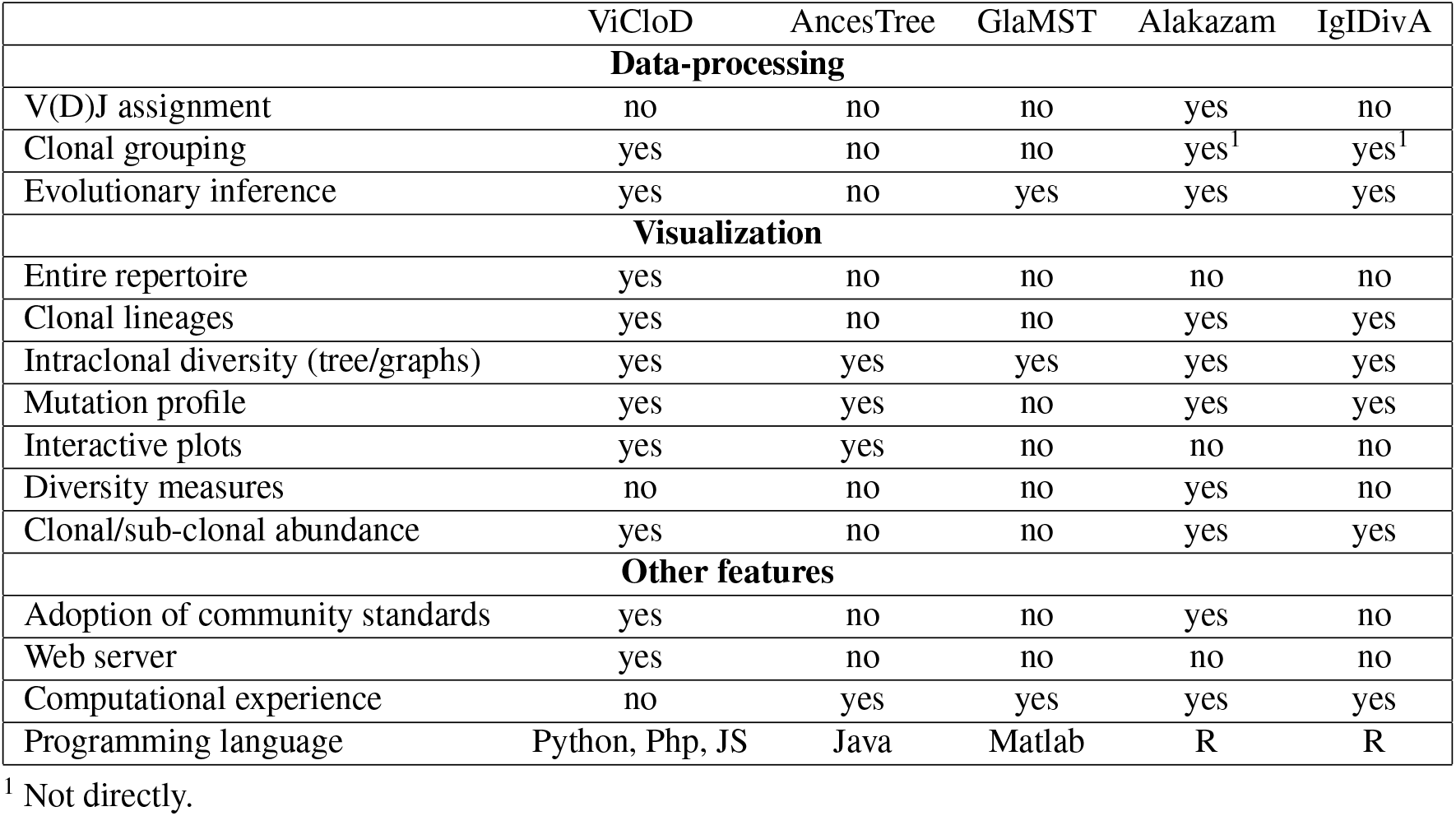
Comparing ViCloD with other clonal lineage visualization tools. We summarized the features of ViCloD and compared them to tools that achieve evolution analysis and visualize it.

AncesTree (*38*) is a stand-alone graphic interface to interactively explore clonal lineage trees. It does not carry out any data processing; instead, it parses the outputs of Dnaml (*55*) and IgPhyML (*56*). AncesTree displays pre-computed trees, the number of nucleotide and amino acid mutations, and multiple sequence alignments. Trees are interactive, and users can visualize the detailed location of each mutation. To explore intraclonal diversity, users need to pre-process several tasks, such as clonal grouping, clone selection and lineage reconstruction. Another stand-alone tool is GLaMST (*39*) that uses the minimum spanning tree algorithm to infer clonal lineage trees from high-throughput sequencing data. GLaMST runs in Matlab and requires input sequences in FASTA format, where the first sequence represents the HAUS (root tree). Like Ancestree, GlaMST does not process the raw data, and the analysis of intraclonal diversity requires computational experience. Furthermore, it does not provide interactive plots or mutational profiles.

Alakazam (*40*) is part of the Immcantation framework. Users need to run Change-O (*33*) to detect clonal groups, and then select a clonal lineage of interest, for example, the most abundant one. The clonal lineage tree inference is performed by dnapars (*55*), which constructs a maximum parsimonious tree of a given clonal lineage. Alakazam provides a set of trees with different layouts, but they are not interactive, and they do not display clonal abundance. An overview of the repertoire composition is not direct, but can be obtained by running other tools from the Immcantation framework such as Sumrep (*36*). However, Alakazam computes several diversity measures, such as the species richness, and the Shannon index, and provides gene usage, and chemical properties of the amino acid sequences, which are not yet available in ViCloD.

IgIDivA (*41*) is a dedicated tool for intraclonal analysis; it first identifies all nucleotide variants (here termed as clonotypes) inside a subclone. Second, it constructs a network/graph centered on the most abundant clonotype rather than a tree from the HAUS. Two clonotypes are connected only if their connection is consistent with the progressive acquisition of SHMs. To give an overview of intraclonal diversity, IgIDivA computes a set of graph network metrics, such as average degree and maximal path length in the network, among others. Since IgIDivA analyzes clonotypes instead of subclones, its outputs are not directly comparable to ViCloD ones. Moreover, it does not adopt community standards input files, making its usage more difficult with V(D)J assignment tools other than IMGT/HighV-QUEST.

## 4 Discussion

High-throughput sequencing analysis has made possible quantitative studies in immunology, where adaptive immune receptor repertoires (AIRR) can contain millions of different receptor gene variants. Individual repertoire analysis is now routinely possible and affordable, resulting in many practical applications in biology and medical research. Several web servers devoted to analyzing AIRR data have been recently reported, most of them for pre-processing, V(D)J gene annotation, clonal grouping, repertoire diversity, and mutation analysis (*26*–*37*). However, only some tools permit the analysis of intraclonal diversities and the investigation of the mutational architecture of an individual repertoire and its evolution (*36*, *38*, *40*, *41*). Most existing tools require some computational skills to manage installations and resolve dependencies. Their usage is then limited to research environments since, most of the time, the clinical environment has software installation constraints. Moreover none of them provide an interactive, versatile and anywhere-access tool, with an entire view of the repertoire composition, where users can easily navigate among the most relevant clonal lineages and explore their evolutionary events.

To fill the gap, we propose ViCloD, an interactive web server for visualizing B cell repertoire composition and analyzing intraclonal diversities. We demonstrated the ViCloD usefulness by analyzing several BCR repertoires from patients with different B-cell malignancies, some known for bearing intraclonal diversification in their IG variable regions. This clinical context was chosen since the B cell population is dominated by a single clonal IGH-VDJ rearrangement (or two in case of biallelic rearrangements), which helps visualize intraclonal diversity due to the large number of available sequences. ViCloD has constructed different B-cell lineage tree topologies and mutation profiles for the four analyzed repertoires. We have observed trees with higher depths and more cumulative mutations in patients with higher intraclonal diversity. These study cases demonstrated how ViCloD could analyze this type of data in detail, helping to visualize their intraclonal diversity.

Our web-server can process and analyze data coming directly from IMGT/HighV-QUEST, the international standard web software for annotating IG repertoire sequence data. However, it is not restricted to IMGT/HighV-QUEST outputs since it adopts the AIRR file format (*45*). In principle, the output of any V(D)J annotation tool can be used since it produces the necessary data http://www.lcqb.upmc.fr/viclod/Format. Although we have applied ViCloD to B-cell malignancies, the server is not limited to analyzing a specific pathology and can be applied to other clinical conditions (*57*).

ViCloD offers several interactive plots, including B-cell lineage trees and mutation profiles. Such tools allow users to interact with their data, which is more complicated with other tools that provide pre-defined outputs (*38*–*41*). ViCloD calculations are performed in real-time, and the results are shown immediately on the screen (or, eventually, a link is sent by email). Therefore, the user can analyze complex data very efficiently and quickly. The web-server is user-friendly and versatile, particularly for analyzing high-throughput sequencing repertoire data, providing several views of BCR repertoire, which can help users to extract important features from their analyses. Moreover, all data and figures are easily downloaded and exported to other applications for further inspection. ViCloD is available to all users without requiring registration or any computational requirement. Users’ data are stored temporarily, but they can be downloaded in a predefined format that allows re-analyzing posteriorly.

Additional features will be implemented in the future, such as (i) intraclonal analysis at the amino acid level, including Physico-chemical properties, (ii) diversity measures, such as the species richness, and the Shannon index, and gene usage plots, (iii) metrics to measure the intraclonal diversity level (*41*), (iv) the possibility of visualizing evolutionary events within a give subclone, (v) comparisons of several repertoires, which will allow users to analyze repertoires from the same individual at various time points or to compare repertoires of different individuals. ViCloD’s flexibility also offers the possibility of further extensions, including the estimation of B cell migration and differentiation (*42*).

## 5 Conclusion

We have produced a new AIRR bioinformatics tool, ViCloD, which can help process, analyze and visualize B cell receptor repertoire clonality and intraclonal diversity in B cell populations. ViCloD provides interactive modules for analyzing and visualizing the repertoire topology, clonal compositions, and clonal lineage architectures. Differently from the most used AIRR-Seq tool, for intraclonal diversity analysis, ViCloD provides a graphical web interface for clinicians, immunologists, and researchers without strong computational backgrounds. It provides interactive plots and allows users to examine thoroughly their data, including sequences, plots, and figures, which are easily exportable. Finally, ViCloD run-time and memory requirements have been minimized in order to be compatible with clinical applications. We believe that, with the growing importance of immune repertoire research, ViCloD will play an important role in identifying specific features within immune repertoires, helping to decipher the high-dimensional complexity of the adaptive immune system.

## Supporting information

S1 Table: Computational time

S2 Table: Properties for the five most abundant clonal lineage.

S1 Figure: BCR repertoire organization.

S2 Figure: Simplifying tree topology.

S3 Figure Clonotype distribution.

