## Supplementary Figures for "ViCloD, an interactive web tool for visualizing B cell repertoires and analyzing intraclonal diversities: application to human B-cell tumors"

### ViCloD, an interactive web tool for visualizing B cell repertoires and analyzing intra-clonal diversities in B-cell tumors. Supplementary File

Lucile Jeusset, Nika Abdollahi, Thibaud Verny, Marine Armand, Anne Langlois De Septenville, Frédéric Davi and Juliana S. Bernardes

February 16, 2023

#### 1 Supplementary Tables

Table S1: **Data processing time for the analysis of four patients.** For each patient, we show the number of reads in the uploaded AIRR file, the number of inferred clonal lineages, and the computational time needed to produce the result pages.

| | Number of inputted reads | Number of clonal lineages | Computational time (seconds) |
|---|---|---|---|
| Patient 1 | 185723 | 173 | 106 |
| Patient 2 | 52220 | 17 | 27 |
| Patient 3 | 25093 | 2 | 15 |
| Patient 4 | 532948 | 33 | 118 |

Table S2: **Top five clonal lineages of a patient with Follicular Lymphoma (Patient 1).** For each clonal lineage, we show its label, abundance in the repertoire (Ab%), the absolute number of analyzed reads, IGHV, IGHJ gene annotations, and CDR3 segment (amino acid sequences). The "functionality" status indicates when sequences are productive and unproductive.

| Clonal lineage | Ab(%) | Number of reads | IGHV | IGHJ | CDR3 | Functionality |
|---|---|---|---|---|---|---|
| C1 | 94.986 | 44258 | IGHV3-23*01 | IGHJ4*02 | VRNRSSHDSQSSYFDY | yes |
| C2 | 0.146 | 68 | IGHV3-23*01 | IGHJ4*02 | AKS*YS*VAVFLL*L | no |
| C3 | 0.086 | 40 | IGHV4-61*02 | IGHJ4*02 | AREVRTFELLSSFDS | yes |
| C4 | 0.075 | 35 | IGHV1-69*06 | IGHJ6*02 | AREGGGPAGMGYYGTDV | yes |
| C5 | 0.071 | 33 | IGHV3-30*03 | IGHJ4*02 | AKDGYGDYGEYFDY | yes |

#### 2 Supplementary Figures

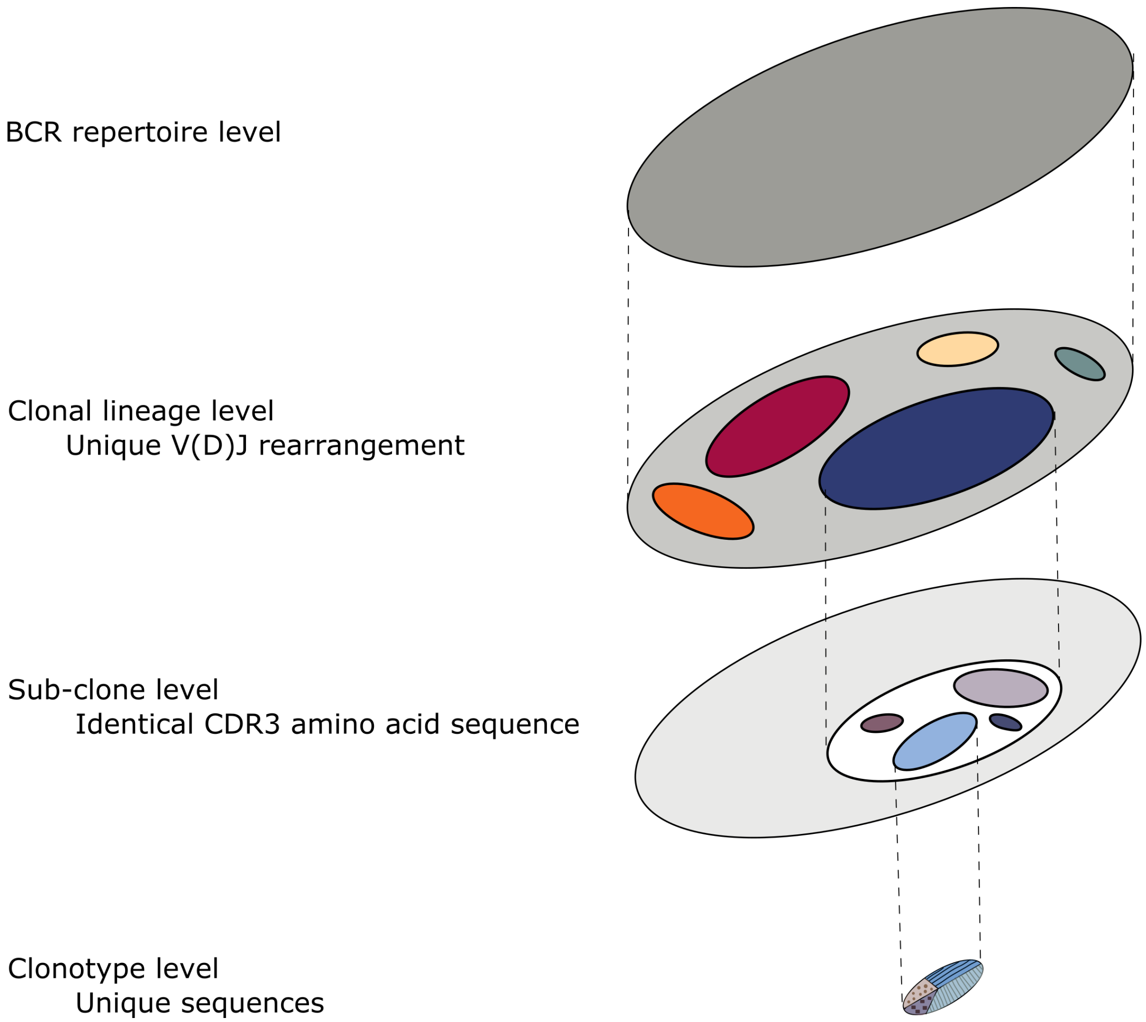


Figure S1: **BCR repertoire organization**. The first level represents the entire set of sequences of a give repertoire. The second level represents B cell lineages. The third level groups sequences with identical CDR3 amino acid content, forming a sub-clone. The fourth level groups identical nucleotide sequences within a given sub-clone, termed as clonotype level.

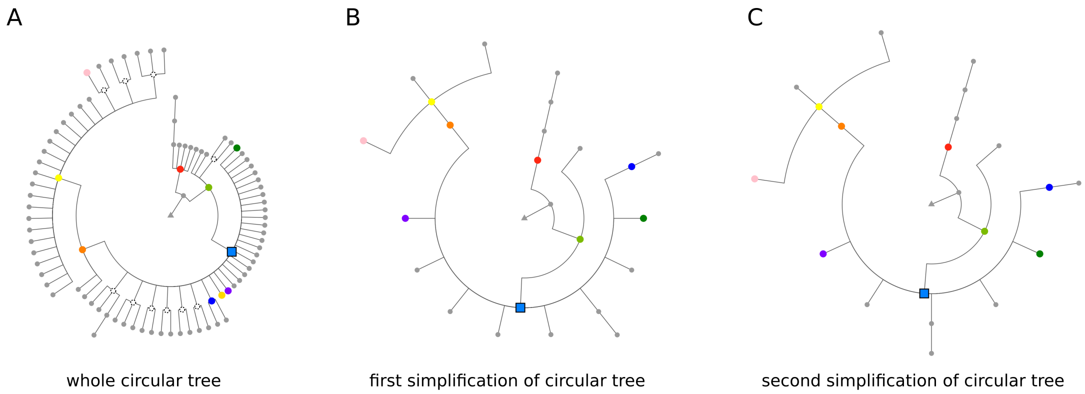


Figure S2: **Simplifying tree topology.** Simplification of B-cell lineage tree (A) of clonal lineage C1 of patient1. The first simplification (B) eliminates the nodes which have no descendants or whose abundance is lower than a certain threshold (fixed at 0.01). The second simplification (C) takes the first 30 nodes of the first simplification and eliminates the nodes with the lowest Hamming distance.

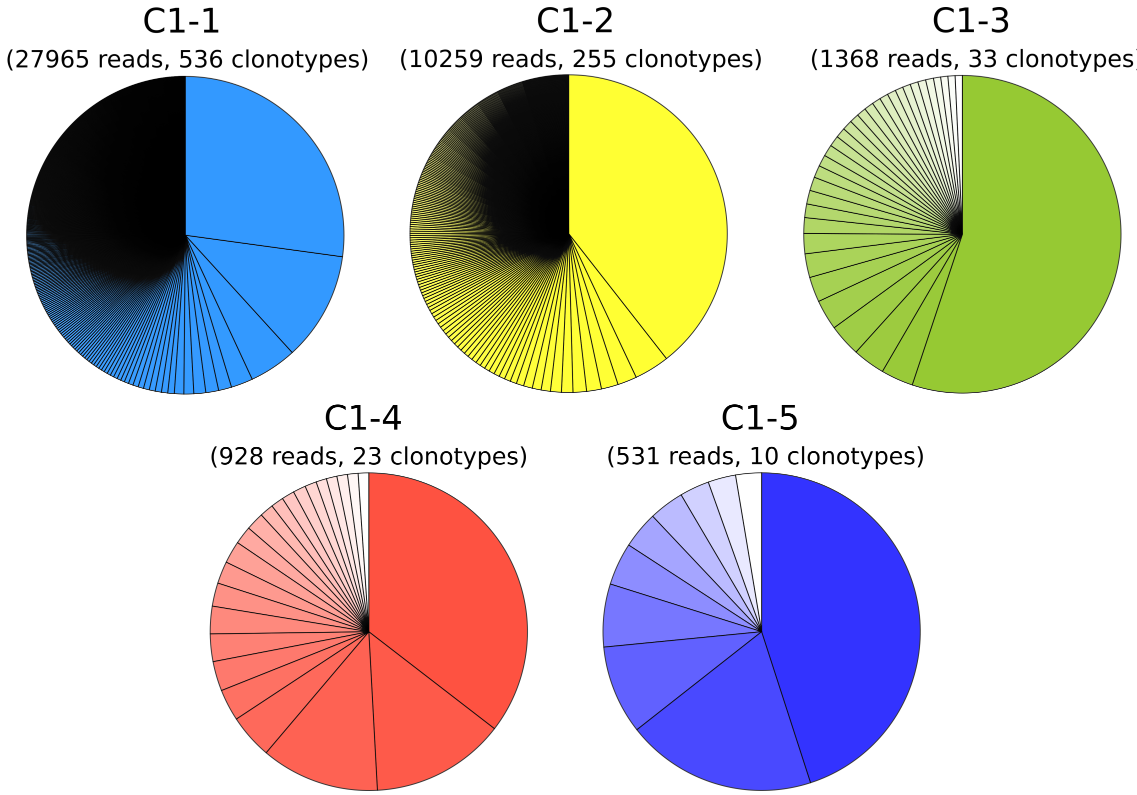


Figure S3: **Clonotype distribution**. Each pie chart represents the proportion of clonotype within the 5 most abundant subclones of the clonal lineage C1. The total number of sequences contained in each subclone is indicated above the pie chart as well as the number of clonotypes contained in this subclone.
